# Whole genome sequencing for drug resistance profile prediction in *Mycobacterium tuberculosis*

**DOI:** 10.1101/401703

**Authors:** Sebastian M. Gygli, Peter M. Keller, Marie Ballif, Nicolas Blöchliger, Rico Hömke, Miriam Reinhard, Chloé Loiseau, Claudia Ritter, Peter Sander, Sonia Borrell, Jimena Collantes Loo, Anchalee Avihingsanon, Joachim Gnokoro, Marcel Yotebieng, Matthias Egger, Sebastien Gagneux, Erik C. Böttger

**Author notes:** Corresponding authors Prof. Erik C. Böttger, MD, University of Zurich, Institute of Medical Microbiology, National Center for Mycobacteria, Gloriastrasse 28/30, 8006 Zurich, Switzerland; Prof. Sebastien Gagneux, PhD, Swiss Tropical and Public Health Institute, Socinstrasse 57, 4002 Basel, Switzerland. equal contribution as first authors (Sebastian M. Gygli, Peter M. Keller). equal contribution as last authors (Matthias Egger, Sebastien Gagneux, Erik C. Böttger).

## Abstract

Whole genome sequencing allows rapid detection of drug-resistant *M. tuberculosis* isolates. However, high-quality data linking quantitative phenotypic drug susceptibility testing (DST) and genomic data have thus far been lacking.

We determined drug resistance profiles of 176 genetically diverse clinical *M. tuberculosis* isolates from Democratic Republic of the Congo, Ivory Coast, Peru, Thailand and Switzerland by quantitative phenotypic DST for 11 antituberculous drugs using the BD BACTEC MGIT 960 system and 7H10 agar dilution to generate a cross-validated phenotypic DST readout. We compared phenotypic drug susceptibility results with predicted drug resistance profiles inferred by whole genome sequencing.

Both phenotypic DST methods identically classified the strains into resistant/susceptible in 73-99% of the cases, depending on the drug. Changes in minimal inhibitory concentrations were readily explained by mutations identified by whole genome sequencing. Using the whole genome sequences we were able to predict quantitative drug resistance levels where wild type and mutant MIC distributions did not overlap. The utility of genome sequences to predict quantitative levels of drug resistance was partially limited due to incompletely understood mechanisms influencing the expression of phenotypic drug resistance. The overall sensitivity and specificity of whole genome-based DST were 86.8% and 94.5%, respectively.

Despite some limitations, whole genome sequencing has high predictive power to infer resistance profiles without the need for time-consuming phenotypic methods.

**One sentence summary:** Whole genome sequencing of clinical *M. tuberculosis* isolates accurately predicts drug resistance profiles and may replace culture-based drug susceptibility testing in the future.

## Introduction

Timely and accurate drug susceptibility testing (DST) of *M. tuberculosis* isolates is vital to prevent the transmission of multidrug-resistant strains (MDR – resistance to rifampicin and isoniazid)[1]. However, the slow growth and stringent biosafety requirements of *M. tuberculosis* make obtaining a full DST profile by culture-based techniques a matter of weeks or months. In addition, culture-based DST is notoriously challenging for several drugs, e.g. pyrazinamide and ethionamide due to poor drug solubility in commonly used culture media [2].

Drug resistance in *M. tuberculosis* is mainly conferred by chromosomal mutations in a few genes [3], making it possible to detect drug resistance by sequencing these genes or probing them by molecular hybridisation [4]. Several commercial tests for the detection of resistance-associated mutations are available, e.g. the GenoType MTBDRplus V2 (Hain Lifescience GmbH, Nehren, DE) [5], the AID TB Resistance Line Probe Assay (AID GmbH, Strassberg, DE) [6]. Line probe assays and the GeneXpert^®^ system (Cepheid, Sunnyvale, CA, USA) are endorsed by the World Health Organisation (WHO) the detection of rifampicin resistance as surrogate marker for MDR [7]. These molecular tests demonstrate high sensitivities for drugs with established target(s) of resistance and for which only a few mutations are responsible for most resistance *in clinico* (e.g. rifampicin, isoniazid) [4]. However, molecular tests show low sensitivity for heteroresistant strains (concomitant presence of wild type (wt) and resistant or multiple different resistant variants in patient isolates), when frequencies of resistant variants drop below 5-50 % [8, 9]. Furthermore, there are no commercially available tests for many drugs currently/prospectively in use (e.g. bedaquiline, delamanid, linezolid, p-aminosalicylic acid).

The past years have seen a wealth of genomic data on drug-resistant *M. tuberculosis* become available [10, 11]. However, phenotypic DST data are lacking for most of the genetic data sets. In addition, DST data are often limited as the strains were classified as susceptible or resistant using only a single drug concentration. There is an urgent need to link genotypic and phenotypic drug resistance readouts to obtain a better understanding of the mechanisms influencing the evolution and spread of drug resistance in *M. tuberculosis*.

WGS of clinical isolates allows for accurate identification of established-resistance-conferring chromosomal mutations [10, 12, 13] and may ensure adequate treatment in days instead of months. We compared whole genome-based drug resistance profiles with two culture-based quantitative DST methods for a total of 11 drugs, including all first-line drugs (rifampicin, isoniazid, ethambutol, pyrazinamide, streptomycin) and an array of second-line drugs (rifabutin, amikacin, kanamycin A, capreomycin, moxifloxacin, ethionamide).

## Material and methods

### *M. tuberculosis* isolates

The initial data-set consisted of 190 *M. tuberculosis* isolates. A subset of 61 strains was used to establish the phenotypic DST methodology. These 61 strains were collected by the Swiss National Center for Mycobacteria between 2004-2015, and represent a broad spectrum in geographic origin and drug resistance profiles [14–16]. We then applied the quantitative DST methodology to 125 clinical isolates from clinics participating in the International Epidemiology Databases to Evaluate AIDS (IeDEA) [17] in Peru, Thailand, Ivory Coast and the Democratic Republic of the Congo (supplementary Table S3). Thirteen strains had to be excluded due to failed WGS (n = 4, failed library preparation due to low DNA quality), irreproducible DST results (n = 1), no growth in the 7H10 agar dilution assay (n = 3), duplication (n = 1), mixed cultures (n = 2, cross-contamination or patient infected with multiple strains) or transmission clusters (n = 2). The final set consisted of 176 strains.

### Phenotypic DST

MGIT 960- and 7H10 agar dilution-based phenotypic DST were performed as described previously [14]. Table 1 lists the epidemiological cut-offs (ECOFF) used [18], supplementary Table S2 the drug concentrations tested with the MGIT 960 and 7H10 agar-dilution assays and Table 2 the genes screened for mutations with WGS. Further details are available in the supplementary materials.

**Table 1:** Epidemiological cutoffs (ECOFF) used for 7H10 agar dilution and MGIT 960 phenotypic DST [14]

| Antibiotic | ECOFF<br>agar<br>dilution<br>(mg/L) | ECOFF<br>MGIT<br>960<br>(mg/L) |
| --- | --- | --- |
| Ethionamide | 1 | 5 |
| Ethambutol | 2 | 5 |
| Capreomycin | 4 | 2.5 |
| Streptomycin | 0.5 | 1 |
| Kanamycin A | 2 | 2 |
| Amikacin | 1 | 1 |
| Moxifloxacin | 0.25 | 0.25 |
| Isoniazid | 0.125 | 0.1 |
| Rifampicin | 0.5 | 1 |
| Rifabutin | 0.0625 | 0.1 |
| Pyrazinamide | NA | 100 |

**Table 2:** List of genes implicated in drug resistance in *M. tuberculosis* which were screened for polymorphisms by WGS. List adapted from [3, 21]

| Drug | Target gene(s) |
| --- | --- |
| Ethionamide | *ethA*, *inhA*, *inhA* promoter |
| Ethambutol | *embB* |
| Capreomycin | *rrs*, *eis* promoter, *tlyA* |
| Streptomycin | *rrs*, *gidB*, *rpsL* |
| Kanamycin A | *rrs*, *eis* promoter |
| Amikacin | *rrs*, *eis* promoter |
| Moxifloxacin | *gyrA* |
| Isoniazid | *katG*, *inhA* promoter |
| Rifampicin/rifabutin | *rpoB* |
| Pyrazinamide | *pncA*, *pncA* promoter |

### Data analysis

The categorical agreement between the MIC determination by MGIT 960 and 7H10 agar dilution was determined based on the ECOFFS (Table 1).

The numerical variation between the two methods was quantified as the geometric standard deviation (SD, given with its standard error) of the ratio MIC MGIT 960/MIC agar dilution, expressed as a number of 2-fold dilutions and denoted by σ. The geometric SD was computed by fitting a log-normal distribution to the ratio MIC MGIT 960/MIC agar dilution as implemented in the R package fitdistrplus (v.1.0-9) [19]. If the data was compatible with σ = 0, the geometric standard deviation could not be estimated and was defined as “not applicable” (NA). The approach is a generalization of the Bland and Altman method [20], taking censoring of the data into account. Strains for which the MGIT 960 MIC and 7H10 agar dilution MIC were both left-censored or both right-censored were excluded since no information on the ratio could be derived.

Goodman and Kruskal’s gamma was used to quantify the rank correlation between the two methods. No correlation could be calculated if the variance for either method was 0 (NA).

Distributions of wt and mutant MICs were analysed qualitatively based on the results of 7H10 agar dilution. We divided the dataset into two groups: drugs for which the MIC distributions of wt and mutant strains did not overlap, and those for which MIC distributions overlapped.

Sensitivities and specificities of WGS-based resistance profile inference were calculated based on the 7H10 agar dilution results for all drugs, except pyrazinamide – for which the MGIT 960 results were used, based on resistance/susceptibility at the WHO-defined critical concentrations and the presence or absence of a putative resistance-associated mutation.

### Defining clinical breakpoints for high/low-level resistance

The therapeutic window of a drug is defined as the maximal serum concentration which is considered safe [21]. Mutations can increase the MIC beyond the therapeutic window and render the drug clinically ineffective. Drugs may have large therapeutic windows beyond the ECOFF. For these, MIC increases caused by mutations may still be within the therapeutic window of a drug: these strains might still be treatable by increasing the drug dose. We analysed the distribution of MICs of mutant strains, and assessed if cut-offs for low-level (within the therapeutic window) and high-level (beyond the therapeutic window) resistance were definable.

### WGS and single nucleotide polymorphism (SNP) calling

WGS and data analysis was performed as previously described [22] and summarised in the supplementary materials. The performance of WGS-based DST greatly depends on the availability of robust markers of resistance. We therefore focussed on a set of high-confidence resistance-associated genes [4, 12] (Table 2).

### Ethics

Local institutional review board or ethics committee approval was obtained at all local study sites. Informed consent was obtained where requested per local regulations. This project was approved by the Swiss Ethics Committee on research involving humans (swissethics, Bern, Switzerland).

## Results

### Agreement between MGIT 960 and 7H10 agar dilution phenotypic DST

Table 3 and Figure 2 summarize the agreement between the semi-quantitative/quantitative MIC determination by MGIT 960 and 7H10 agar dilution in terms of classifying strains as resistant or susceptible according to ECOFFs (Table 1). Agreement was high for all drugs, except ethambutol (see below). For most drugs, the MGIT 960-based MICs were higher than the 7H10 agar dilution-based MICs. MICs obtained using the two methods were within 1-2 two-fold dilution steps of each other. The classifications into resistant or susceptible demonstrated high rank correlations (Table 3 and Figure 2), except for capreomycin (supplementary Figure S4) due to few resistant strains included in the study.

**Table 3:**
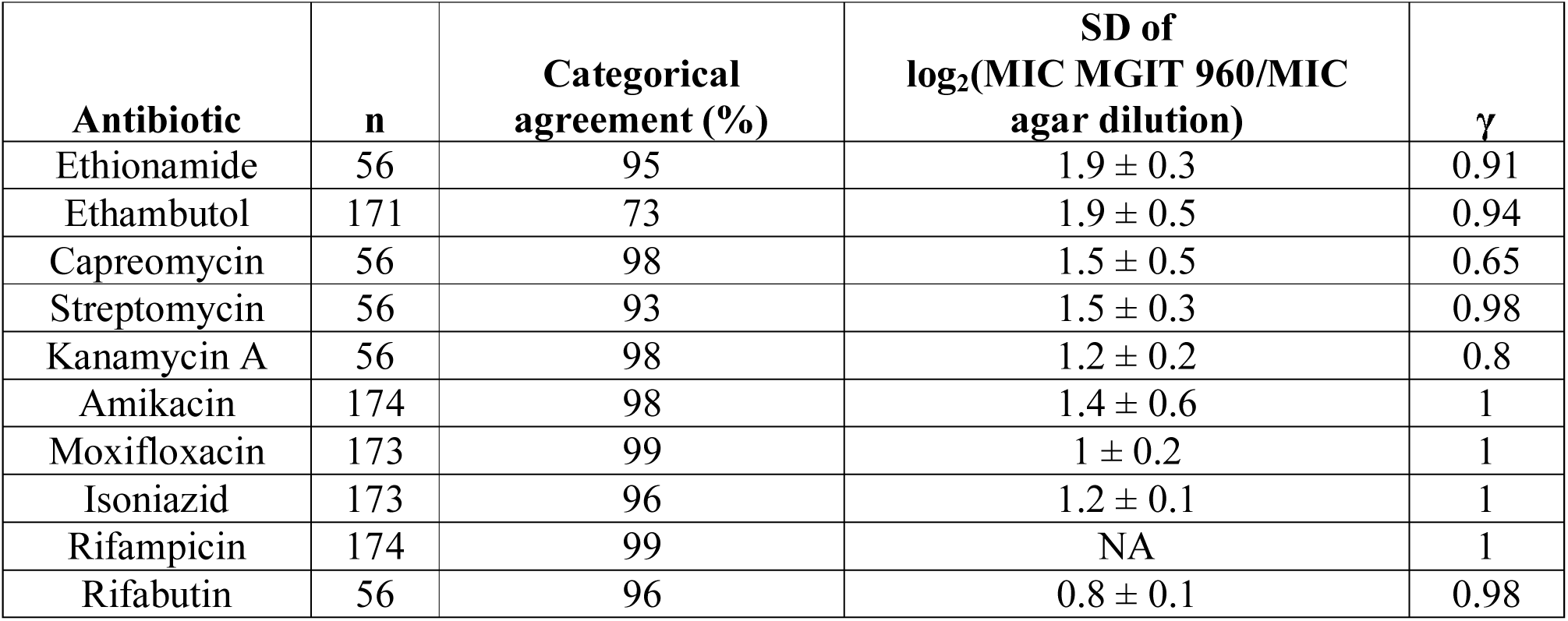
Summary statistics of the method agreement between 7H10 agar dilution- and MGIT 960-based phenotypic DST for all drugs assayed in this study

| Antibiotic | n | Categorical agreement (%) | SD of $\log_2(\text{MIC MGIT 960/MIC agar dilution})$ | $\gamma$ |
| --- | --- | --- | --- | --- |
| Ethionamide | 56 | 95 | $1.9 \pm 0.3$ | 0.91 |
| Ethambutol | 171 | 73 | $1.9 \pm 0.5$ | 0.94 |
| Capreomycin | 56 | 98 | $1.5 \pm 0.5$ | 0.65 |
| Streptomycin | 56 | 93 | $1.5 \pm 0.3$ | 0.98 |
| Kanamycin A | 56 | 98 | $1.2 \pm 0.2$ | 0.8 |
| Amikacin | 174 | 98 | $1.4 \pm 0.6$ | 1 |
| Moxifloxacin | 173 | 99 | $1 \pm 0.2$ | 1 |
| Isoniazid | 173 | 96 | $1.2 \pm 0.1$ | 1 |
| Rifampicin | 174 | 99 | NA | 1 |
| Rifabutin | 56 | 96 | $0.8 \pm 0.1$ | 0.98 |

### WGS and *in silico* resistance profile prediction

A total of 176 WGS with a median coverage of 67.6X (interquartile range (IQR) = 37.48) were obtained. Median mapping percentage and percentage of genome covered were 98.7% (IQR = 0.94) and 99.4% (IQR = 0.4), respectively. All major *M. tuberculosis* lineages, except lineage 7, were represented in the study (L1 = 6, L2 = 36, L3 = 11, L4 = 123, L5 = 1, L6 = 1). The strains showed a range of drug resistance profiles (Figure 1). Based on the set of analysed genes (Table 2), 25 strains were predicted to be fully susceptible against all assayed drugs, 59 strains were mono-/poly-resistant, 91 strains demonstrated MDR phenotypes and two strains were extensively drug resistant (XDR: isoniazid, rifampicin, fluoroquinolone and aminoglycoside resistant).

**Figure 1:**
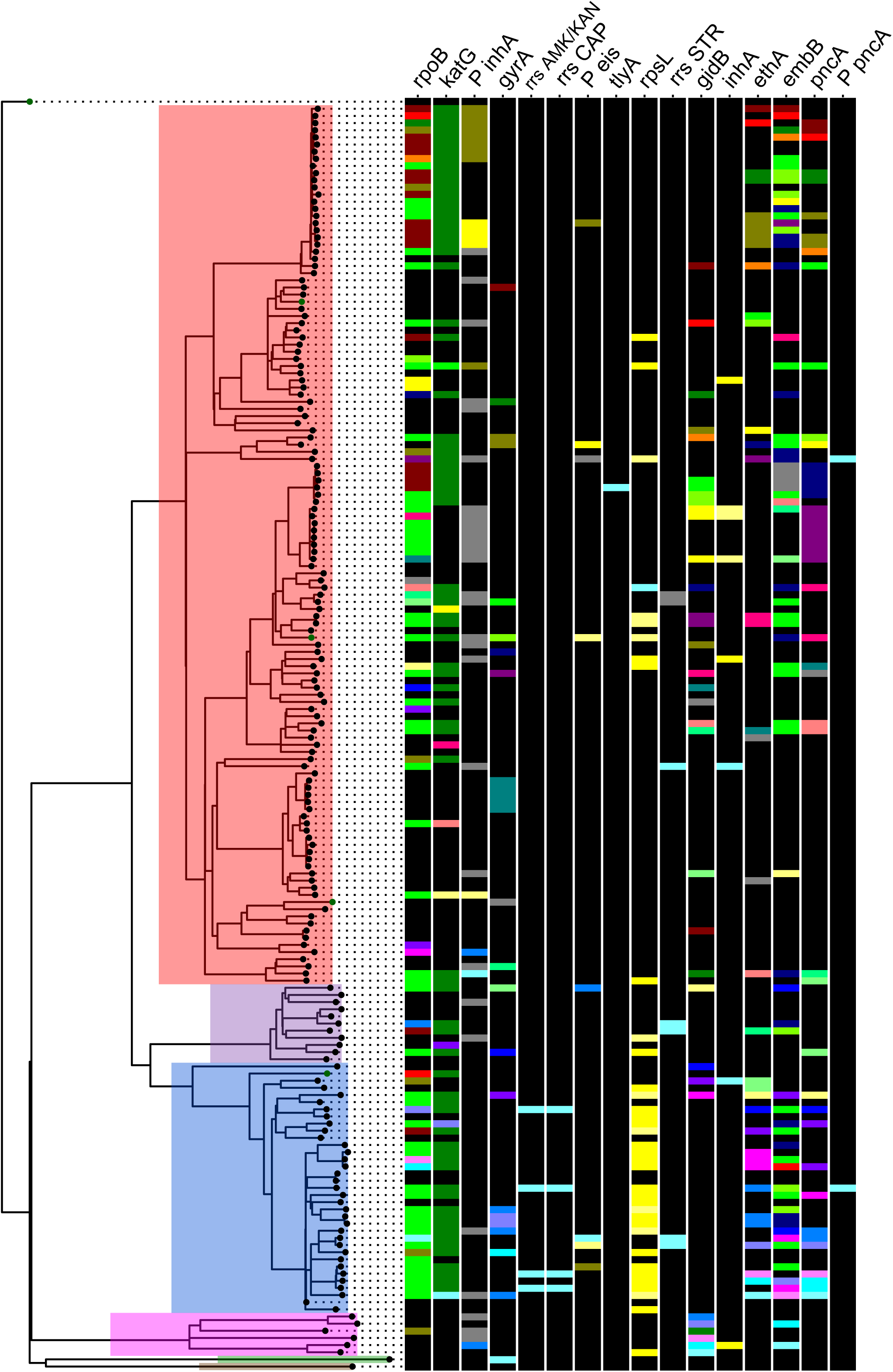
Maximum likelihood phylogeny of 176 *M. tuberculosis* strains based on 20510 variable positions. Reference strains labeled with green tip labels. Main lineages are highlighted as follows: red L4, purple L3, blue L2, pink L1, green L6, brown L5. Scale bar indicates number of substitutions per site. Phylogeny rooted on *M. canettii*. Colored bars indicate resistance mutations per gene and within a distinct column (gene) each colored bar represents a distinct mutation. Black bars indicate no mutation, i.e. wt.

**Figure 2:**
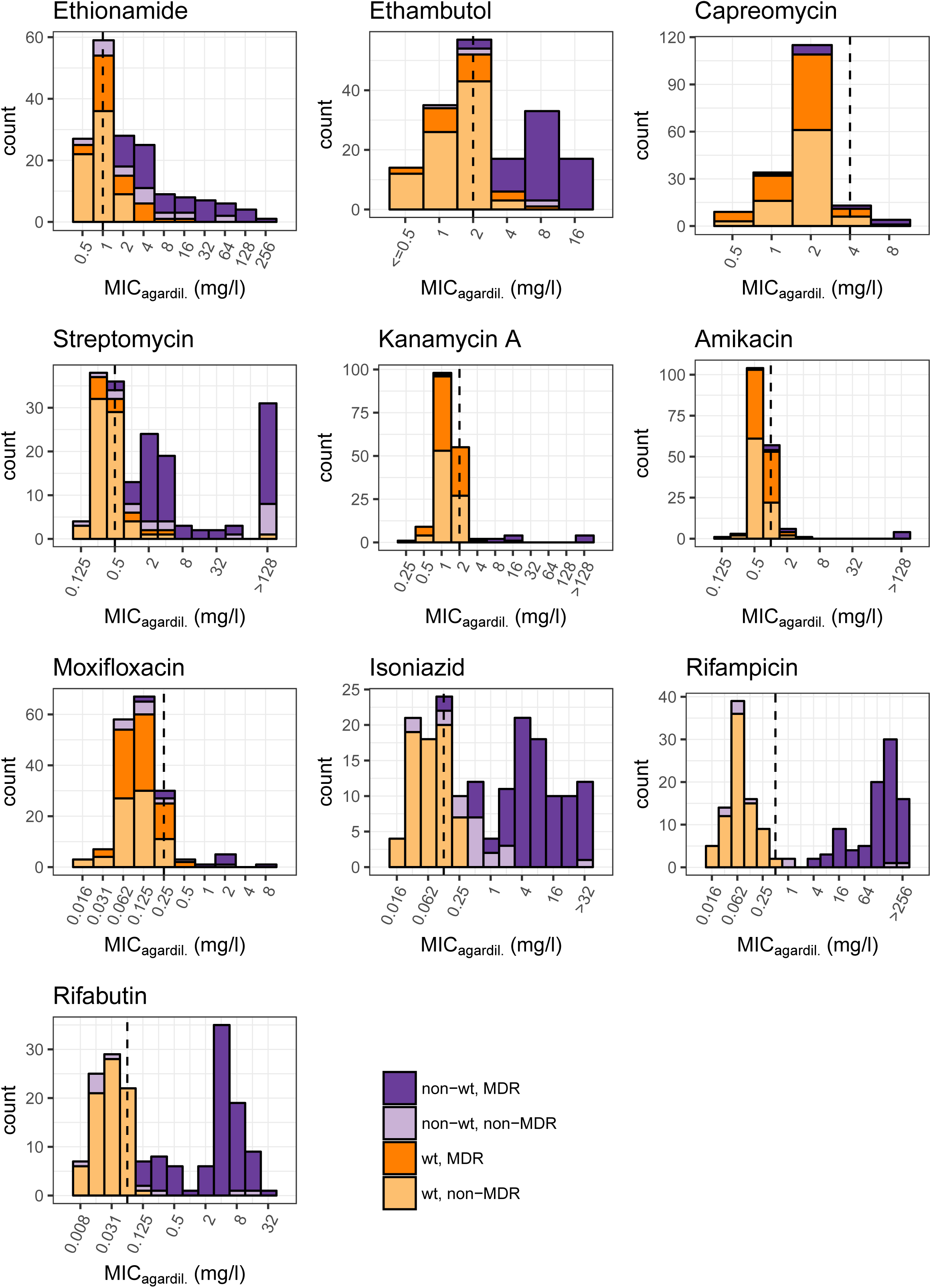
Histograms of MICs (7H10 agar dilution) for all drugs assayed in this study

### Drug resistance profile prediction by WGS vs. phenotypic DST

After exclusion of known phylogenetic markers not involved in resistance, WGS-based prediction of drug resistance using a defined set of target genes (Table 2) was highly congruent with the categorical classification based on the phenotypic DST for most drugs (Table 3 and Figure 2). Based on the *in silico* resistance prediction, the MICs of mutant and wt strains frequently followed a Gaussian distribution. However, the same resistance marker may conferre different MICs in different strains (supplementary Figures S1C, S2C, S3C, S8C, S9C, S10C). In some cases, the increase in the MIC conferred by a certain resistance mutation fell within the distribution the of wt MIC (e.g. for *gidB, eis* promotor mutations, supplementary Figures S3C, S6C).

### Distinct wt and mutant MIC distributions

MIC distributions of wt and mutant strains were well separated for rifampicin, rifabutin, isoniazid, kanamycin A, amikacin, capreomycin, streptomycin and pyrazinamide, indicating that the resistance markers used had a high positive predictive power (88.9% overall positive predictive power of resistance markers). For streptomycin, two strains harboured no mutations in the target genes, yet demonstrated high-level phenotypic resistance (supplementary Figure S3C).

### Overlapping wt and mutant MIC distributions

MIC distributions of wt and mutant strains overlapped for ethambutol, moxifloxacin and ethionamide. For ethambutol and ethionamide, overlapping MIC distributions of wt and mutant strains were associated with a large number of polymorphisms in resistance-conferring genes (ethambutol resistance: 22 polymorphisms in *embB*, ethionamide resistance: 28 in *ethA*, 3 in *inhA*, 6 in *inhA* promoter). Solubility issues with ethionamide led to quantitative differences in MGIT 960 vs. 7H10 agar dilution-based DST (Table 3, Figure 3). The overlap in MIC distributions between wt and strains carrying an *embB* mutation was reduced by adjusting the critical concentration for ethambutol resistance from 5 mg/L to 2.5 mg/L (MGIT 960). However, there was variability in the MICs for the same mutation (e.g. MIC EmbB M306I/V in 7H10 agar dilution: 4-16 mg/L –supplementary Figure S2C). Moxifloxacin resistance was rare (n = 9, MGIT 960, critical concentration 0.25 mg/L) and MIC distributions of mutant strains partially overlapped with those of wt. Sensitivity of the genome-based moxifloxacin resistance prediction was 80.0% (Table 4).

**Table 4:** Sensitivity and specificity of the genome-based drug resistance profile prediction using the 7H10 agar dilution-based categorical classification as the gold standard for all drugs except pyrazinamide, for which the MGIT 960 categorical classification was used

| Drug | Sensitivity (%) | Specificity (%) |
| --- | --- | --- |
| Ethionamide | 75.0 | 92.9 |
| Ethambutol | 89.6 | 94.2 |
| Capreomycin | 75.0 | 94 |
| Streptomycin | 68.0 | 92.1 |
| Kanamycin A | 83.3 | 98.8 |
| Amikacin | 63.6 | 96.9 |
| Moxifloxacin | 80.0 | 90.2 |
| Isoniazid | 93.6 | 96.8 |
| Rifampicin | 100 | 94.0 |
| Rifabutin | 98.9 | 94.0 |
| Pyrazinamide | 80.8 | 88.9 |

**Figure 3:**
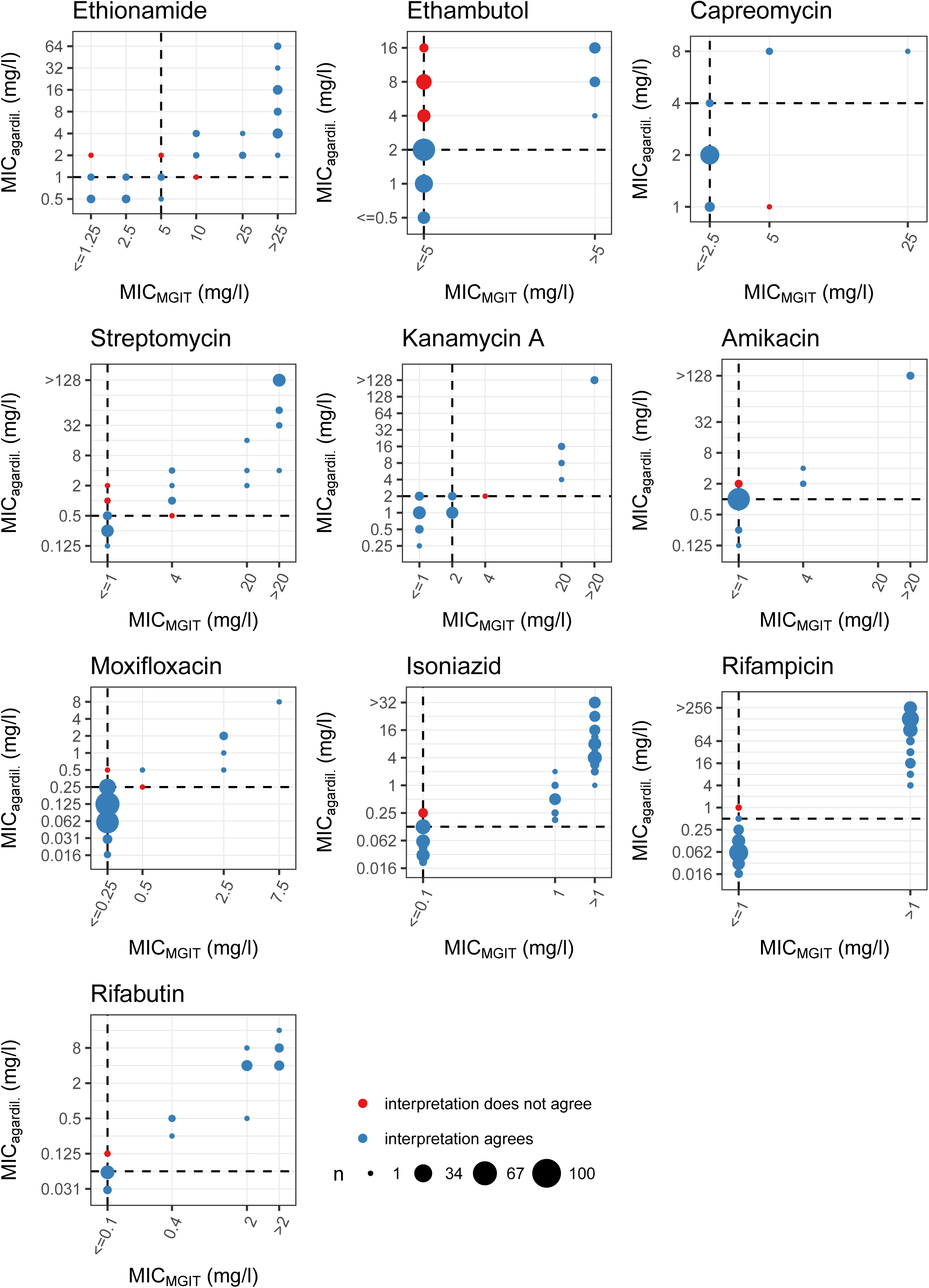
Method agreement between phenotypic DST performed with MGIT 960 and 7H10 agar dilution represented as Bland-Altman plots for all drugs tested in this study.

### Defining high-/low-level clinical breakpoint concentrations

#### Isoniazid

Mutations in the promoter of *inhA* conferred low-level resistance <1 mg/L (7H10 agar dilution), compared to strains harbouring mutations in *katG* or combinations of *inhA* promoter and *katG* mutations which demonstrated MIC levels ranging from >1 mg/L to >32 mg/L in 7H10 agar dilution (supplementary Figure S8C). Defining clinical breakpoint concentrations (CBC) for low-(≤1 mg/L for MGIT 960/7H10 agar dilution) and high-level (>1 mg/L MGIT 960/7H10 agar dilution) isoniazid resistance is warranted.

### Rifampicin/Rifabutin

Most mutations in *rpoB* increased the MIC for rifamycins beyond the therapeutic window (peak serum concentration 10 mg/L [21, 23]). However, some rare *rpoB* mutations (e.g. RpoB L452P, H445Y – supplementary Figure 9C) demonstrated MICs within the therapeutic window. Defining low- and high-level CBC may thus be justified.

For rifampicin, CBC were ≤4/2 mg/L for MGIT 960/7H10 agar dilution and >4/2 mg/L for MGIT 960/7H10 agar dilution, respectively.

For rifabutin, our data suggests CBC for low- and high-level resistance of ≤0.4/0.25 or 0.5 mg/L for MGIT 960/7H10 agar dilution and >0.4/0.25 or 0.5 mg/L for MGIT 960/7H10 agar dilution, respectively.

Mutations in *rpoB* conferring resistance to rifampicin and rifabutin showed highly correlated increases (Figure 4) of MICs beyond the therapeutic window for most *rpoB* mutations (Figure 3 and supplementary Figure S9C & S10C), indicating that both drugs are rendered clinically ineffective and cannot substitute each other.

**Figure 4:**
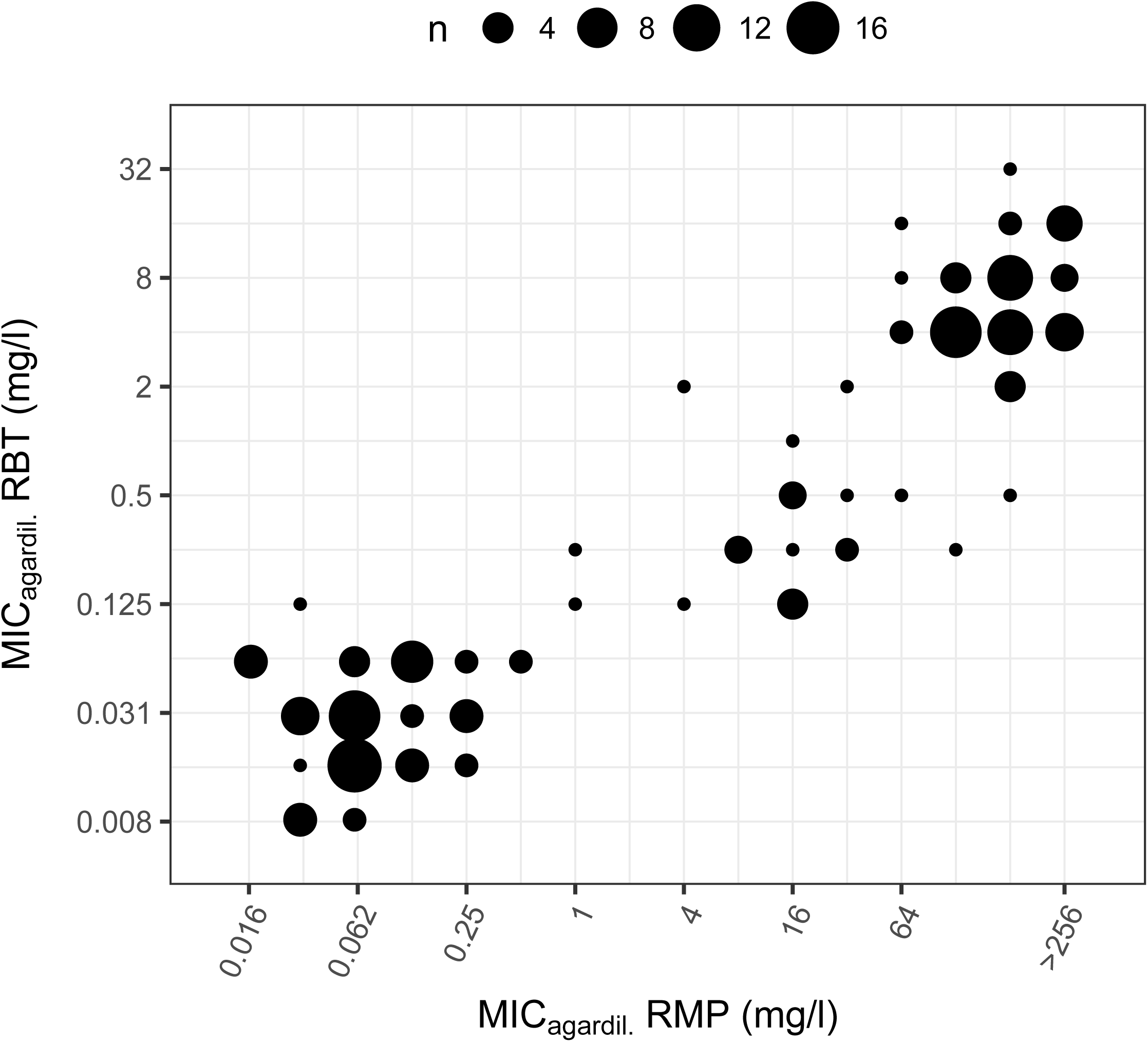
Correlation between 7H10 agar dilution MICs for rifampicin and rifabutin

### Amikacin

Few strains had mutations in the regions of *rrs* relevant for amikacin resistance or the *eis* promoter (n=12). Mutations in *rrs* were associated with high-level (>128 mg/L in 7H10 agar dilution) and mutations in the promoter region of *eis* with low-level level (2-4 mg/L in 7H10 agar dilution) amikacin resistance. Given the peak serum concentrations of amikacin (20-40 mg/L [21]), a CBC for low-(≤ 4 mg/L for MGIT 960/7H10 agar dilution) and high-level (4 mg/L for MGIT 960/7H10 agar dilution) amikacin resistance may be warranted.

### Streptomycin

Certain mutations lead to MICs well beyond the peak serum concentrations [21] of streptomycin (e.g. RpsL K43R, MIC 7H10 agar dilution >128 mg/L, supplementary Figure S3C). On the other hand, *gidB* mutations increase the MIC only moderately (MIC 7H10 agar dilution 1-4 mg/L, supplementary Figure 3C). Mutational combinations in *gidB, rrs, rpsL* were common and produced a range of different MICs. However, there were mutations that systematically lead to MICs beyond the therapeutic window, e.g. RpsL K43R. Defining low-level (MGIT 960 ≤4 mg/L, 7H10 agar dilution ≤4-8 mg/L) and high-level CBC for streptomycin resistance (MGIT 960 >4 mg/L, 7H10 agar dilution >4-8 mg/L) is warranted.

## Discussion

We compared quantitative phenotypic DST with *in silico* or genomic resistance profile prediction inferred from WGS using 176 clinical *M. tuberculosis* isolates.

The results of MGIT 960 and 7H10 agar dilution-based phenotypic DST methods were highly correlated and suitable to separate susceptible from resistant variants. After exclusion of known phylogenetic markers, genome-based resistance profile prediction displayed high sensitivity and specificity for detecting resistance. Based on phenotypic DST results and WGS, we were able to define CBC for high- and low-level resistance for isoniazid, rifampicin, streptomycin and amikacin. Defining such breakpoints is important for preserving efficacious drugs for treatment of resistant *M. tuberculosis* variants.

Our data suggest that the current WHO-defined critical concentration for phenotypic DST of ethambutol by MGIT 960 (5 mg/L) is too high and may misclassify strains as susceptible when compared to the 7H10 agar dilution-based classification. Given the low peak serum concentrations for ethambutol, this may lead to mistreatment due to presumed ethambutol susceptibility. After adjusting the critical concentration to 2.5 mg/L for MGIT 960, we observed a strong improvement of the categorical agreement between MGIT 960- and 7H10 agar dilution-based classification.

The mutations identified by WGS had a high predictive power to classify strains as resistant. However, the predictive power depends on a number of factors. For instance, the increase in MIC conferred by an identical resistance mutation can vary greatly in different strains (e.g. EmbB M306I/V, RpsL K88R). Such variation is clinically relevant if there is a significant overlap between the MICs of mutant and wt strains, as was the case for ethionamide, ethambutol and streptomycin (e.g. *gidB*) resistance mutations. Furthermore, it is difficult to classify strains as resistant or susceptible if the MIC increase lies within the therapeutic window of a drug. The overlap between MICs of mutant and wt strains is confounded by the fact that we only screened for mutations in genes which had previously been associated with drug resistance. We might thus have missed possible resistance-confering mutations in other genes. Additionally, WGS will always produce distributions of coverages which in term will inevitably lead to certain regions in the genome suffering from low coverage, preventing the detection of mutations. The inability to call mutations due to low coverage will therefore lead to false negatives, reducing sensitivity. Furthermore, the strain genetic background [24], non-mutational mechanisms (e.g. modulation of gene expression) [25], as well as drug efflux mechanisms [26] may contribute to the variability in increase of the MIC conferred by resistance mutations.

The predictive power of mutations in target genes also depends on removing phylogenetic markers not involved in resistance. Separating phylogenetic from resistance-associated markers works well for essential (highly conserved) genes such as *rpoB, rpsL, rrs* but is problematic in non-essential genes involved in the conversion of prodrugs into their active forms like *pncA* (pyrazinamide), *ethA* (ethionamide) or in genes that generally exhibit higher numbers of polymorphisms e.g. *embB*. Of note, the *embABC* operon is highly polymorphic, harbouring more polymorphisms than expected by chance (mutations in *embABC* operon = 81, expected = 44.8, p = 9.174e-07, binomial test). Mutations conferring ethambutol resistance [27] will therefore inevitably evolve in the presence of phylogenetic SNPs and may interact epistatically to produce the variability in MICs we observed for wt strains and for the most common ethambutol resistance markers *embB* M306I/V. The *embAB*C operon is involved in the biosynthesis of decaprenylphosphoryl-β-d-arabinose, which is an integral component of the mycobacterial cell wall. The cell envelope interacts with the host immune system and the high levels of diversity of these genes might be the product of diversifying selection due to host immune pressure. The influence of polymorphisms in the *embABC* operon on MICs in general is supported by the observation that sub-inhibitory concentrations of ethambutol lower the MICs for isoniazid, rifampicin and streptomycin [28]. Even small changes in activity of the decaprenylphosphoryl-β-d-arabinose biosynthetic and utilisation pathway might thus alter cell wall permeability and influence MICs of several drugs.

Similarly, in the case of streptomycin resistance, the RpsL substitution K88R exhibited a range in MICs from low to high-level resistance making it difficult to judge the susceptibility of a strain harbouring this mutation based on the genotype. Streptomycin was the first effective antituberculous drug discovered [29] and has been used for decades. The long-term use has produced complex resistance profiles with multiple streptomycin resistance mutations (e.g. in *gidB, rpsL, rrs*) occurring concomitantly, producing wide ranges of MICs. Furthermore, many streptomycin resistant strains displayed MDR/XDR phenotypes. Streptomycin resistance mutations are frequently found in backgrounds which have mutations in genes affecting the information pathway (DNA -> RNA -> proteins) – e.g. *gyrA* (DNA gyrase), *rpoB* (DNA-dependent RNA polymerase), *rrs* (ribosomal RNA). The simultaneous presence of multiple resistance mutations may alter the adaptive landscape [30, 31]. In addition, non-mutational processes (e.g. alteration of gene expression) may compensate for fitness costs of drug resistance and at the same time alter the MIC for the drug [25]. This has not been demonstrated for streptomycin resistance in *M. tuberculosis*, but it seems possible that compensation of fitness costs in MDR phenotypes might alter the MIC for streptomycin [30], considering that streptomycin is not part of the current standard treatment regimen and selection for high-level streptomycin resistance is relaxed.

In conclusion, we demonstrate that MGIT 960 and 7H10 agar dilution-based phenotypic DST provide highly congruent classifications of strains into resistant or susceptible. WGS has high predictive power to infer resistance profiles without the need for time-consuming phenotypic methods. Limitations due to overlapping distributions of wt and mutant MICs, varying MICs for the same resistance mutations in different strains, presence of phylogenetic markers in resistance-associated genes and rare resistance markers with low frequencies will likely be resolved by on-going large-scale projects (e.g. ReSeqTB [32]) combining phenotypic DST with WGS of thousands of *M. tuberculosis* isolates. Our findings, together with those of on-going studies will pave the way for the replacement of phenotypic DST with drug resistance profile prediction based on WGS in the coming years.

## Acknowledgements

We would like to thank Alexandra Mushegian for critically reading the manuscript and improving the writing. Whole genome analysis was performed at sciCORE (http://scicore.unibas.ch/) scientific computing core facility at University of Basel.

## References

1. World Health Organization. Treatment of tuberculosis: Guidelines [Internet]. 4th Editio. Treat. Tuberc. Guidel. 2010. Available from: http://www.ncbi.nlm.nih.gov/books/NBK138741/#ch2.s3.

2. Domínguez J, Boettger EC, Cirillo D, Cobelens F, Eisenach KD, Gagneux S, Hillemann D, Horsburgh R, Molina-Moya B, Niemann S, Tortoli E, Whitelaw A, Lange C, TBNET, RESIST-TB networks. Clinical implications of molecular drug resistance testing for Mycobacterium tuberculosis: a TBNET/RESIST-TB consensus statement. Int. J. Tuberc. Lung Dis. [Internet] 2016; 20: 24–42Available from: http://www.ncbi.nlm.nih.gov/pubmed/26688526.

3. Gygli SM, Borrell S, Trauner A, Gagneux S. Antimicrobial resistance in Mycobacterium tuberculosis: mechanistic and evolutionary perspectives. FEMS Microbiol. Rev. [Internet] 2017;: 1–20 Available from: https://academic.oup.com/femsre/article-lookup/doi/10.1093/femsre/fux011.

4. Deggim-Messmer V, Bloemberg G V., Ritter C, Voit A, Hömke R, Keller PM, Böttger EC. Diagnostic Molecular Mycobacteriology in Regions With Low Tuberculosis Endemicity: Combining Real-time PCR Assays for Detection of Multiple Mycobacterial Pathogens With Line Probe Assays for Identification of Resistance Mutations. EBioMedicine [Internet] The Authors; 2016; 9: 228–237Available from: http://dx.doi.org/10.1016/j.ebiom.2016.06.016.

5. Nathavitharana RR, Hillemann D, Schumacher SG, Schlueter B, Ismail N, Omar SV, Sikhondze W, Havumaki J, Valli E, Boehme C, Denkinger CM. Multicenter noninferiority evaluation of hain GenoType MTBDRplus Version 2 and Nipro NTM+MDRTB line probe assays for detection of rifampin and isoniazid resistance. J. Clin. Microbiol. 2016; 54: 1624–1630.

6. Ritter C, Lucke K, Sirgel FA, Warren RW, van Helden PD, Bottger EC, Bloemberg G V. Evaluation of the AID TB Resistance Line Probe Assay for Rapid Detection of Genetic Alterations Associated with Drug Resistance in Mycobacterium tuberculosis Strains. J. Clin. Microbiol. [Internet] 2014; 52: 940–946 Available from: http://jcm.asm.org/cgi/doi/10.1128/JCM.02597-13.

7. WHO. Automated Real-Time Nucleic Acid Amplification Technology for Rapid and Simultaneous Detection of Tuberculosis and Rifampicin Resistance: Xpert MTB/RIF Assay for the Diagnosis of Pulmonary and Extrapulmonary TB in Adults and Children: Policy Update [Internet]. Autom. Real-Time Nucleic Acid Amplif. Technol. Rapid Simultaneous Detect. Tuberc. Rifampicin Resist. Xpert MTB/RIF Assay Diagnosis Pulm. Extrapulm. TB Adults Child. Policy Updat. 2013.Available from: http://www.ncbi.nlm.nih.gov/pubmed/25473701.

8. Engström A. Fighting an old disease with modern tools: characteristics and molecular detection methods of drug-resistant Mycobacterium tuberculosis. Infect. Dis. (London, England) [Internet] 2015; 4235: 1–17Available from: http://www.ncbi.nlm.nih.gov/pubmed/26167849.

9. Streicher EM, Bergval I, Dheda K, Böttger EC, Gey Van Pittius NC, Bosman M, Coetzee G, Anthony RM, Van Helden PD, Victor TC, Warren RM. Mycobacterium tuberculosis population structure determines the outcome of genetics-based second-line drug resistance testing. Antimicrob. Agents Chemother. 2012; 56: 2420–2427.

10. Coll F, McNerney R, Preston MD, Guerra-Assunção JA, Warry A, Hill-Cawthorne G, Mallard K, Nair M, Miranda A, Alves A, Perdigão J, Viveiros M, Portugal I, Hasan Z, Hasan R, Glynn JR, Martin N, Pain A, Clark TG. Rapid determination of anti-tuberculosis drug resistance from whole-genome sequences. Genome Med. [Internet] 2015; 7: 51 Available from: http://genomemedicine.com/content/7/1/51.

11. Walker TM, Kohl TA, Omar S V, Hedge J, Del C, Elias O, Bradley P, Iqbal Z, Feuerriegel S, Niehaus KE, Wilson DJ, Clifton DA, Kapatai G, Ip CLC, Bowden R, Drobniewski FA, Allix-béguec C, Gaudin C, Parkhill J, Diel R, Supply P, Crook DW, Smith EG, Walker AS, Ismail N, Niemann S. Whole-genome sequencing for prediction of Mycobacterium tuberculosis drug susceptibility and resistanceL: a retrospective cohort study. 2015; 3099: 1–10.

12. Shea J, Halse TA, Lapierre P, Shudt M, Kohlerschmidt D, Van Roey P, Limberger R, Taylor J, Escuyer V, Musser KA. Comprehensive Whole-Genome Sequencing and Reporting of Drug Resistance Profiles on Clinical Cases of Mycobacterium tuberculosis in New York State. J. Clin. Microbiol. [Internet] 2017; 55: 1871–1882 Available from: http://jcm.asm.org/content/55/6/1871.full.pdf%0Ahttp://ovidsp.ovid.com/ovidweb.cgi?T=JS&PAGE=reference&D=emexa&NEWS=N&AN=616382615.

13. Colman RE, Anderson J, Lemmer D, Lehmkuhl E, Georghiou SB, Heaton H, Wiggins K, Gillece JD, Schupp JM, Catanzaro DG, Crudu V, Cohen T, Rodwell TC, Engelthaler DM. Rapid Drug Susceptibility Testing of Drug Resistant Mycobacterium tuberculosis Directly from Clinical Samples using Amplicon Sequencing: A Proof of Concept Study. J. Clin. Microbiol. [Internet] 2016; 54: JCM.00535-16Available from: http://jcm.asm.org/lookup/doi/10.1128/JCM.00535-16%5Cn https://github.com/TGenNorth/SMOR.

14. Springer B, Lucke K, Calligaris-Maibach R, Ritter C, Böttger EC. Quantitative drug susceptibility testing of Mycobacterium tuberculosis by use of MGIT 960 and EpiCenter instrumentation. J. Clin. Microbiol. 2009; 47: 1773–1780.

15. Stucki D, Ballif M, Egger M, Furrer H, Altpeter E, Battegay M, Droz S, Bruderer T, Coscolla M, Borrell S, Zürcher K, Janssens J-P, Calmy A, Mazza Stalder J, Jaton K, Rieder HL, Pfyffer GE, Siegrist HH, Hoffmann M, Fehr J, Dolina M, Frei R, Schrenzel J, Böttger EC, Gagneux S, Fenner L. Standard Genotyping Overestimates Transmission of Mycobacterium tuberculosis among Immigrants in a Low-Incidence Country. Carroll KC, editor. J. Clin. Microbiol. [Internet] 2016; 54: 1862–1870Available from: http://jcm.asm.org/lookup/doi/10.1128/JCM.00126-16.

16. Bloemberg G V., Keller PM, Stucki D, Trauner A, Borrell S, Latshang T, Coscolla M, Rothe T, Hömke R, Ritter C, Feldmann J, Schulthess B, Gagneux S, Böttger EC. Acquired Resistance to Bedaquiline and Delamanid in Therapy for Tuberculosis. N. Engl. J. Med. [Internet] 2015; 373: 1986–1988Available from: http://www.ncbi.nlm.nih.gov/pubmed/26559594.

17. Egger M, Ekouevi DK, Williams C, Lyamuya RE, Mukumbi H, Braitstein P, Hartwell T, Graber C, Chi BH, Boulle A, Dabis F, Wools-Kaloustian K. Cohort Profile: The international epidemiological databases to evaluate AIDS (IeDEA) in sub-Saharan Africa. Int. J. Epidemiol. [Internet] 2012; 41: 1256–1264Available from: https://academic.oup.com/ije/article-lookup/doi/10.1093/ije/dyr080.

18. World Health Organization. Technical Report on critical concentrations for drug susceptibility testing of medicines used in the treatment of drug-resistant tuberculosis [Internet]. 2018.Available from: http://www.who.int/iris/handle/10665/260470.

19. Delignette-Muller ML, Dutang C. fitdistrplus: an R package for fitting distributions. J. Stat. Softw. 2015; 64: 1–34.

20. Martin Bland J, Altman D. STATISTICAL METHODS FOR ASSESSING AGREEMENT BETWEEN TWO METHODS OF CLINICAL MEASUREMENT. Lancet [Internet] 1986; 327: 307–310Available from: http://www.sciencedirect.com/science/article/pii/S0140673686908378.

21. Böttger EC. The ins and outs of Mycobacterium tuberculosis drug susceptibility testing. Clin. Microbiol. Infect. [Internet] 2011; 17: 1128–1134 Available from: http://linkinghub.elsevier.com/retrieve/pii/S1198743X14629396.

22. Ghielmetti G, Coscolla M, Ruetten M, Friedel U, Loiseau C, Feldmann J, Steinmetz HW, Stucki D, Gagneux S. Tuberculosis in Swiss captive Asian elephants: microevolution of Mycobacterium tuberculosis characterized by multilocus variable-number tandem-repeat analysis and whole-genome sequencing. Sci. Rep. [Internet] 2017; 7: 14647 Available from: http://www.nature.com/articles/s41598-017-15278-9.

23. Sekaggya-Wiltshire C, von Braun A, Lamorde M, Ledergerber B, Buzibye A, Henning L, Musaazi J, Gutteck U, Denti P, de Kock M, Jetter A, Byakika-Kibwika P, Eberhard N, Matovu J, Joloba M, Muller D, Manabe YC, Kamya MR, Corti N, Kambugu A, Castelnuovo B, Fehr JS. Delayed Sputum Conversion in TB-HIV Co-Infected Patients with Low Isoniazid and Rifampicin Concentrations. Clin. Infect. Dis. [Internet] 2018; Available from: https://academic.oup.com/cid/advance-article/doi/10.1093/cid/ciy179/4919550.

24. Fenner L, Egger M, Bodmer T, Altpeter E, Zwahlen M, Jaton K, Pfyffer GE, Borrell S, Dubuis O, Bruderer T, Siegrist HH, Furrer H, Calmy A, Fehr J, Stalder JM, Ninet B, Bottger EC, Gagneux S. Effect of Mutation and Genetic Background on Drug Resistance in Mycobacterium tuberculosis. Antimicrob. Agents Chemother. [Internet] 2012 [cited 2013 Aug 9]; 56: 3047–3053 Available from: http://www.pubmedcentral.nih.gov/articlerender.fcgi?artid=3370767&tool=pmcentrez&rendertype=abstract.

25. Freihofer P, Akbergenov R, Teo Y, Juskeviciene R, Andersson DI, Böttger EC. Nonmutational compensation of the fitness cost of antibiotic resistance in mycobacteria by overexpression of tlyA rRNA methylase. RNA [Internet] 2016; 22: 1836–1843Available from: https://www.ncbi.nlm.nih.gov/pubmed/27698071.

26. da Silva PEA, Machado D, Ramos D, Couto I, Von Groll A, Viveiros M. Efflux Pumps in Mycobacteria: Antimicrobial Resistance, Physiological Functions, and Role in Pathogenicity. In: Li X-Z, Elkins CA, Zgurskaya HI, editors. Efflux-Mediated Antimicrob. Resist. Bact. [Internet] Cham: Springer International Publishing; 2016. p. 527–559 Available from: http://link.springer.com/10.1007/978-3-319-39658-3.

27. Safi H, Lingaraju S, Amin A, Kim S, Jones M, Holmes M, McNeil M, Peterson SN, Chatterjee D, Fleischmann R, Alland D. Evolution of high-level ethambutol-resistant tuberculosis through interacting mutations in decaprenylphosphoryl-β-D-Arabinose biosynthetic and utilization pathway genes. Nat. Genet. [Internet] Nature Publishing Group; 2013; 45: 1190–1197 Available from: http://dx.doi.org/10.1038/ng.2743.

28. Jagannath C, Reddy VM, Gangadharam PRJ. Enhancement of drug susceptibility of multi-drug resistant strains of Mycobacterium tuberculosis by ethambutol and dimethyl sulphoxide. J. Antimicrob. Chemother. [Internet] 1995; 35: 381–390Available from: https://academic.oup.com/jac/article-lookup/doi/10.1093/jac/35.3.381.

29. Schatz A, Bugle E, Waksman SA. Streptomycin, a Substance Exhibiting Antibiotic Activity Against Gram-Positive and Gram-Negative Bacteria. Exp. Biol. Med. 1944; 55: 66–69.

30. Moura de Sousa J, Balbontín R, Durão P, Gordo I. Multidrug-resistant bacteria compensate for the epistasis between resistances. de Visser A, editor. PLOS Biol. [Internet] 2017; 15: e2001741 Available from: http://dx.plos.org/10.1371/journal.pbio.2001741.

31. Borrell S, Teo Y, Giardina F, Streicher EM, Klopper M, Feldmann J, Muller B, Victor TC, Gagneux S. Epistasis between antibiotic resistance mutations drives the evolution of extensively drug-resistant tuberculosis. Evol. Med. Public Heal. [Internet] 2013 [cited 2013 May 30]; 2013: 65–74 Available from: http://emph.oxfordjournals.org/cgi/doi/10.1093/emph/eot003.

32. Starks AM, Avilés E, Cirillo DM, Denkinger CM, Dolinger DL, Emerson C, Gallarda J, Hanna D, Kim PS, Liwski R, Miotto P, Schito M, Zignol M. Collaborative Effort for a Centralized Worldwide Tuberculosis Relational Sequencing Data Platform: Figure 1. Clin. Infect. Dis. [Internet] 2015; 61: S141–S146 Available from: https://academic.oup.com/cid/article-lookup/doi/10.1093/cid/civ610.

